# Postural Stability During Illusory Self-Motion - Interactions of Vision and Touch

**DOI:** 10.1101/2023.11.29.569272

**Authors:** Yingjia Yu, Avijit Bakshi, James R. Lackner, Ashton Graybiel

**Affiliations:** Spatial Orientation Laboratory, Brandeis University, Waltham, MA 02454

**Author notes:** Correspondence: Avijit Bakshi Phone Address: MS 033, Brandeis University, 415 South Street, Waltham, MA 02454.

## Abstract

The role of vision in stabilizing balance has been studied exhaustively. Other studies have shown that non-supportive light touch of the fingertip with a surface also can significantly stabilize postural balance. We have studied how vision and cutaneous information jointly affect balance. We used a head-mounted display to simulate a virtual room that rotated about a vertical axis centered with the standing subject’s z-axis. Subjects viewing the display’s rotational displacement soon experienced self-motion and displacement. We assessed how the moving visual input destabilized posture and how it interacted with touch cues that stabilized posture. A novel result is how balance is influenced by the onset of visual motion and the illusion of self-rotation. We discovered that motion perceptions are coupled with stochastic aspects of balance. Changes in the perception of types of motion – none, environment-, and self-rotation – distinctively influence metrics that encode for the stochasticity of balance and do not influence those that filter the stochasticity out. We reconfirmed the significant effects of touch in stabilizing balance and discovered how it interacts with the visual perception of motion. We also found lingering effects of past motion perception, which keep influencing the stochasticity of balance even when visual motion is long stopped. Our findings provide insights into multisensory interaction effects in postural balance and suggest novel future research directions.

## Introduction

Human postural behavior involves two major factors: postural orientation and postural equilibrium (Horak 2006). Studying postural balance involves measuring the extent of the center of pressure or center of mass excursions (Wang, Skubic et al. 2010). Postural balance tasks are usually done with subjects standing on a force plate that directly measures forces exerted in anterior-posterior and medial-lateral directions, reflecting changes in the center of pressure during postural sway.

Studies of human postural balance indicate a complex system relating information from sensory organs and muscular activities. Such studies have demonstrated that visual information, vestibular and proprioceptive inputs, and somatosensory information, mainly from feet and ankles, are required to maintain an upright posture.

### Vision modulates balance

Vision is an important sensory channel for aiding balance control during quiet standing. Vision involves both optical flow and central (also called focal) and peripheral (also called ambient) (Uchiyama and Demura 2009). Gibson used optic flow to describe how visual stimulation plays a role when a person moves in extra-personal space. He later showed its important contribution to the sense of self-motion (Gibson 1950). The central field focuses on object motion perception and recognition, while the peripheral field is sensitive to environmental movements. The perception of self-motion and the direction of displacement are attributed to optical flow in the peripheral visual field, which assists in postural control and balance stabilization (Gibson 1950, Horiuchi, Sato et al. 2017). Motion perception can result from movement of objects in the environment (afferent) or from movements of the eyes, body, or head (efferent) (Kapoula and Lê 2006). Retinal slip, caused by displacement relative to the eyeball, can elicit optokinetic eye movements and feedback for attenuating sway (Guerraz and Bronstein 2008).

Peripheral vision is important for maintaining stable quiet stance (Gaerlan, 2010). The visual stimulation of the peripheral visual field decreases postural sway in the direction of the observed visual stimulus (Berencsi, Ishihara et al. 2005). Guerraz & Bronstein have concluded “peripheral dominance often reported in either visual stabilization of spontaneous body sway or visually induced body sway is more likely due to the size of stimulated field manipulated than to functional specialization of the peripheral vision for postural control” (Guerraz and Bronstein 2008). Visual modulation of postural stability is a function of multiple parameters including object size, jitteriness, binocular disparity, visual acuity, depth of field, spatial frequency, visual motion, and so on. Thus, postural stability increases with the richness of the visual environment (Guerraz and Bronstein 2008).

Subjects undergoing visually induced self-motion (vection) from optic flow show large individual differences in vection strength. The extent to which vision causes such variations has been evaluated by measuring sway differences in center of pressure with visual motion exposure or with eyes closed, and with both seated and standing postures tested (Apthorp, Nagle et al. 2014, Apthorp and Palmisano 2014). Three out of four measures of postural control taken during quiet stance were predicted by measures that varied individually during seated exposure to visual motion and created apparent self-motion (vection).

It is well known that visual motion can destabilize posture balance by inducing illusions of self-motion. However, it is not known whether the different phases of visual motion perception, in which external allocentric rotation is perceived first and egocentric self-motion later, affect posture differentially. Postural control is a complex system requiring multisensory integration. Most studies do not emphasize the complexity of postural balance and the extent to which cognitive processing is involved (Horak 2006). The vestibular system provides a real-time estimate of self-motion in six dimensions, three axes of translation and three axes of rotation, through the otoliths organs and semicircular canals (Cullen 2019). In postural balance research, apparent self-motion is typically generated by optic flow from visual stimulation, although tactile motion stimulation of the soles of the feet can induce illusory self-rotation and displacement (Lackner and DiZio 2013). It is unclear to what extent the vestibular system spatial information generated by postural sway is affected by visual motion and what the interaction would be with somatosensory touch at parietal cortical areas involving multisensory convergence.

### Touch stabilizes balance

Previous studies have shown that non-mechanically supportive fingertip contact with a stationary surface during one legged stance increases postural stability (Holden, Ventura et al. 1994) and fingertip contact levels that cannot physically counteract body sway attenuate tandem Romberg stance sway (Jeka and Lackner 1994).

In touch stabilization studies subjects stood on a force plate measuring the active ground forces generated by body sway in terms of medial-lateral and anterior-posterior center of foot pressure coordinates (Jeka and Lackner 1994). Finger touch forces were measured and found to correlate to the center of pressure sway in the AP and ML directions with a 300 ms delay. Mean sway amplitude (MSA) and CP sway mean power frequency (MPF) were measured as a function of time and indicated that CP MSA in both light and force touch contact was smaller than in no-touch conditions with and without vision, supporting that finger contact reduces body sway. In particular, light touch was more effective than vision in supporting postural balance in the tandem Romberg stance (Jeka and Lackner 1994).

Studying the difference between fingertip contact with a stationary surface versus a moving surface (Jeka, Schöner et al. 1997) showed that light touch contact of the fingertip with a moving surface led to whole-body entrainment of body sway with the touch bar frequency. The touch cue was too small to provide mechanical coupling. Study of the minimum requirement of touch force and the contact surface texture suggested that high rigidity was important in eliciting maximum postural stabilization (Lackner, Rabin et al. 2001), and light touch around 40g was enough to attenuate postural sway.

Induced destabilization is affected by fingertip touch contact. The vibration of the Achilles tendons of a standing subject causes loss of postural balance (Eklund 1972). Lackner et al. tested whether fingertip contact would override the destabilizing vibration of subjects’ leg muscles under eyes-closed conditions (Lackner, Rabin et al. 2000). The peroneus longus and brevis tendons were stimulated by vibration for finger touch and no-touch conditions. Without finger contact, vibration increased the mean sway amplitude of CP of the body and head compared with no touch – no vibration condition. When touch was provided, no significant difference was seen whether vibration was present or not. The fingertip contact was sufficient to suppress the instability caused by tendon vibration that would otherwise give rise to an illusory motion of forward falling. When touch and no-touch conditions were compared under peroneus tendon vibration with on and off vibration periods, results indicated that the longer vibration was presented, the less disturbance subjects experienced regardless of on and off periods when the finger touch cue was provided. Similarly, subjects were more continuously disrupted by vibration regardless of on and off periods when no touch was provided. During touch conditions, removing the vibration exhibited a mirror image rebound, indicating that fingertip contact allowed an active compensation that is anticipatory to counteract the vibration. However, unidirectional shifts were observed in no-touch conditions, indicating that compensations in no-touch conditions were not present. With less than 1 N fingertip touch on a rigid touch bar, there were significantly minor postural disruptions, compensations occurring within 500 ms of vibration onset, and mirror image rebounds on vibration removal (Lackner, Rabin et al. 2000). The effect was stronger with longer periods of vibrations, showing that the nervous system could anticipate the consequences of vibration that causes destabilization and compensate through the stabilization of fingertip contact with a stable and rigid surface to suppress the induced proprioceptive and motor signals in leg muscles.

Biceps branchii vibrations on the arm with light fingertip contact was destabilizing, resulting in greater postural sway amplitudes (Rabin, DiZio et al. 2008). In particular, precision fingertip contact remained intact under vibration, but the vibration created illusory inversion and extension of the fingertip contact arm, destabilizing postural balance. In comparison between touch with arm vibration and touch without arm vibration, the finding that it destabilized more when touch was provided than without was an indication that the increased postural sway amplitudes resulted from the distortion of afferent and efferent information regarding arm configuration (Rabin, DiZio et al. 2008). This was because biceps vibration provided false proprioceptive feedback about arm configuration, and incorrect elbow proprioception about shear forces resulted in an inappropriate posture correction leading to larger sway amplitudes. This finding shows the importance of accurate proprioceptive information about arm configuration during fingertip feedback in postural control tasks.

Individuals lacking vestibular function soon lose balance when they stand heel-to-toe and close their eyes; they fall within seconds in this situation (Lackner, DiZio et al. 1999). Vestibular loss subjects with light touch contact of the right index finger with a rigid bar were more stable than normal subjects without touch during quiet stance in the dark (Lackner, DiZio et al. 1999), demonstrating that the light touch cue could be more effective than vestibular input in minimizing postural sway when no vision was provided. This indicated that within the complex multisensory system, the inclusion of additional somatosensory and proprioceptive information from the touch cue was sufficient to compensate for the loss of vestibular input.

In those experiments, when vision was allowed, it provided exteroceptive information about body position or movement relative to the external environment. In postural control tasks, somatosensory – proprioceptive information is provided through the ankles and feet, which relies largely on a support surface that is held stable. Visual proprioception plays in the form of optic flow, which specifies the movement of the head relative to the environment (Lee and Aronson 1974). Visual motion of the environment presented while subjects are stationary shows that when visual information is misleading, healthy subjects unconsciously try to correct their posture, which leads to either excessive sway or loss of balance (Lee and Lishman 1975, Lee 1977).

### Virtual reality affects perception and balance

VR systems typically employ portable head-mounted displays (HMD) with large fields of view (FOV) updated contingent on head tracking. By adding depth perception and blocking external visual information, VR is a useful tool for investigating the role of vision and visual movement on balance performance (Soltani 2019, Soltani and Andrade 2021).

Subjects immersed in a rotating virtual environment with a vertical axis of rotation will after a delay experience an illusion of self-motion around the yaw axis. This visually induced illusory percept, a.k.a. vection, results from an optic flow field that mimics the visual stimulation of a moving observer (Gibson 1950, Dichgans, Brandt et al. 1978, Lee 1980, Bruder, Steinicke et al. 2011). Vection onset is sensitive to observers’ intent or instructions - to attentively follow the moving background stimulus yields a more delayed vection than suppressing the optokinetic reflex by staring at a fixation pattern in the foreground or even by an inattentive hazy gaze (Becker, Raab et al. 2002, Kitazaki and Sato 2003). The role of higher-level cognitive factors is also born out by the observation that vection is more pronounced when stimulated by ecologically believable, naturalistically rich moving visual stimulation (Schulte-Pelkum, Riecke et al. 2003, Richards, Mulavara et al. 2004, Riecke and Schulte-Pelkum 2006). For example, a naturalistic 3D display or textured room display fosters vection illusion faster, more intensely and with less drop-out than when the stimulus is bland, moving geometrical patterns like polka dots or alternating stripes of an optokinetic drum (Wann and Rushton 1994). Neurophysiological correlates of self-motion illusion have been premised (Hettinger 2002) - a deactivation of early visual cortex and amplitude reductions in visual-evoked potentials (Kleinschmidt, Thilo et al. 2002); greater activations of the right MT+, precuneus, areas located bilaterally along the dorsal part of the intraparietal sulcus and along the left posterior intraparietal sulcus, depth of the left superior frontal sulcus, over the ventral part of the left anterior cingulate, in the depth of the right central sulcus and in the caudate nucleus/putamen, suggesting that the illusory percept of self-motion is correlated with the activation of a network of areas, ranging from motion-specific areas to regions involved in visuo-vestibular integration, visual imagery, decision making, and introspection (Kovács, Raabe et al. 2007).

Vection onset latencies are smaller when participants can uphold their logical credence that their “motion” is “viable” versus when they are cognitively aware that their self-motion is “impossible” (Lepecq, Giannopulu et al. 1995, Palmisano and Chan 2004, Wright, DiZio et al. 2006). For example, a three-dimensional virtual environment may not increase body sway, but optokinetic virtual scenes alter postural control. In a VR setup, optic flow at the eyes may inform the CNS that the surroundings are moving, whereas the somatosensory system says that the body is stationary. Integration in the CNS can either ignore or prioritize conflicting inputs from different sensory modalities (Nashner and Peters 1990, Gaerlan 2010) to accomplish postural control, and it can switch into multiple levels of sensory illusions of vection for either head, torso, and lower body, entire body, or the environmental surfaces touching the body.

In the present study, we create an interaction between visual motion using a well-structured virtual reality motion environment and non-mechanically supportive fingertip contact with a rigid surface in a quiet stance postural balance task. Our goal is to gain insights into the dynamics and complexity of multisensory integration in human postural balance. We investigate how vision and touch interact in the maintenance of postural balance under conflicting conditions with visual motion-induced apparent self-motion and haptic contact with an external surface. For example, will vision dominate and the touched surfaces be experienced as moving with the subject, or will it cause the subject to feel stationary depending on when touch contact is made? Two alternatives will be investigated: as visual motion destabilizes posture, will touch that ordinarily stabilizes posture reduce the postural sway created by vision motion, or would touch provide no additional help, and would self-motion disturb finger touch control. A change in visual motion from no motion-to-motion onset, and from motion-to-motion offset will also be investigated to see the changes in subjects’ center of pressure, weight fraction distribution between the legs, and finger touch force, to gain more understanding of postural control in reaction to potential falling.

## Materials and Methods

### Apparatus and measures

There were three main measuring devices used:

#### Foot force plates

Subjects were required to stand, in quiet stance, on an AMTI (Advanced Mechanical Technology, Inc.) dual force plate throughout the experiment, except for rest periods. Changes in center of foot pressure (CP) on the left and right foot against time were measured separately for medial-lateral (CP_ML_) and anterior-posterior (CP_AP_) coordinates from their 3 force components and torque components. The weight fractions (WF) under the left and right foot were computed from the vertical force under each foot.

#### Touch force plate

A separate AMTI force plate was used to record finger contact forces in all touch trials. The touch contact plate had a smooth square metal surface (225 cm^2^) attached horizontally to a vertical metal stand, parallel to the subjects’ sagittal plane and approximately at the subjects’ waist height. The fingertip touch force magnitude was recorded from the touch plate and CP was computed.

#### Head Mounted Display

An HTC Vive^TM^ Head Mounted Display (HMD), rendering a VR scene at a resolution of 1080 x 1200 and a refresh rate of 90 Hz, was worn throughout the experiment. The HMD had binocular display units with a 110-degree horizontal and vertical field of view. The HMD presented a well-structured environment simulating a living room and shown throughout the experiment. The room had four-walls, ceiling and floor with virtual inner dimensions of width x depth x height = 4.7m x 4.7m x 3m. The room scene was built with *Unity^TM^* software, and the interior was highly textured with doors, windows, sofa, tea table, plant in a pot, and other interior decoration (wall painting, flower vase, etc.). In every trial, half of the time the room was stationary, and the other half it was rotating at 60 degrees per second. The axis of room rotation coincided with the z-axis of the standing subjects. Depending on trial conditions, the room would either start stationary or start with rotation. Room location was adjusted separately for each subject wearing the HMD and standing on the force plate to ensure they seemed to be at the center of the room.

#### Timestamp of perceived self-motion

A smartphone was used during each trial to record the progression of time on the computer screen and the subject’s verbal reporting of self-motion. The experimenter later obtained the exact timestamps of the verbal reports for induced motion from the recordings.

#### Conditions

There were in total 24 trials, divided into two sets of twelve: 12 of the trials presented clockwise room rotation, and the other 12 displayed counterclockwise room rotation. Two independent variables were tested in the experiment:

- Independent variable 1: *touch* or *no touch* of the right index finger
- Independent variable 2: *Start with motion, followed by no motion*, or *Start with no motion, followed by motion*.

Thus, in total, there were 4 difference conditions, all counterbalanced across subjects.

#### Subjects

Fourteen subjects were recruited from Brandeis students and staff, aged between 18-55, 8 females and 6 males. All subjects were in good health and were screened for motion sickness from 3D Virtual Reality exposure. Only one subject was left-handed. All protocols were approved by the Brandeis Institutional Review Board.

#### Procedure

After signing an informed consent form, subjects were shown the apparatus used in the experiment and familiarized with the concept of self-motion by experiencing it. They wore the HMD while sitting on a chair, and viewed room rotation in the HMD and reported verbally when they began to feel self-rotation. In the experimental trials, subjects stood on the feet force plate in quiet stance, without shoes, with their two feet parallel and aligned to the edges of guiding tapes taped placed on the dual foot force plate.

Subjects were asked to close their eyes between trials and open their eyes throughout the trial. Before beginning each trial, subjects were told what the trial condition would be. Subject hand positions were either by their sides relaxed (no touch condition) or using their right index finger to touch the touch force plate on their right side (touch condition). Foot position was checked before the start of every trial. The subject then opened his/her eyes and said “ready” to start the trial. After the “ready” signal was given by the subject, the experimenter responded by saying “collecting” and started the data collection. After the virtual room motion started, the subject reported “moving” when they first started to feel self-motion, which was recorded on the computer and synced with the data collection. A second experimenter served as a “spotter” to aid the subject should loss of balance occur. When each trial ended, the experimenter instructed the subject to relax and close their eyes. Questions regarding dizziness level, sleepiness level, and fatigue level were rated on a 0-10 Likert scale (0 being their state when they entered the laboratory, and 10 being the worst they ever felt). The subjects’ motion perception in relation to the touch plate, foot force plate, and body were asked after certain trials. There were 24 trials in total, each lasting 40 seconds, of which 20 sec was with stationary VR scene and 20 with the rotating VR scene. To avoid fatigue and to allow the experimenter to change rotation direction in the HMD, a 5-minute rest occurred in the middle of the experiment after 12 trials. The total time taken for a complete experiment was approximately 1 hour. All data were collected in real-time, recorded, and saved.

#### Filtering

The CP data (computation described below) and the raw force components data from both force plates were filtered by a 5th-order 5 Hz lowpass Butterworth filter (the *butter* function in MATLAB).

#### Data analysis

*CP in AP, ML, and Net.* CP in AP and ML coordinates were computed from

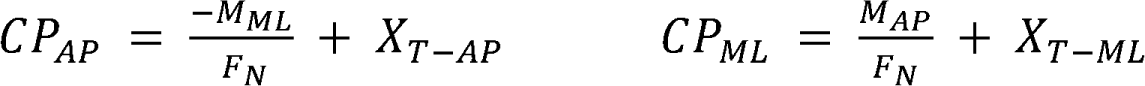

F_N_ represents force applied by subjects normal to the force plate. M represents torque and was taken from both AP and ML directions. X represents offset where origin was centered in the middle of the two feet. The net Center of pressure (CP_net_) was computed from the left and right foot forces and CP.

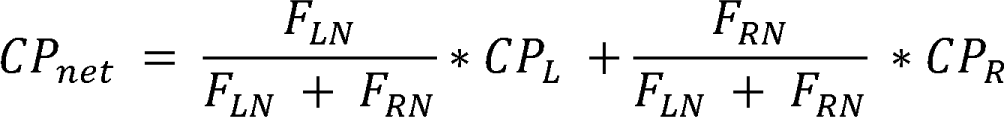

#### Weight Fraction (WF)

The WF was determined from the vertical component (z) of the force (F) measured in the left and right plate using the following equation:

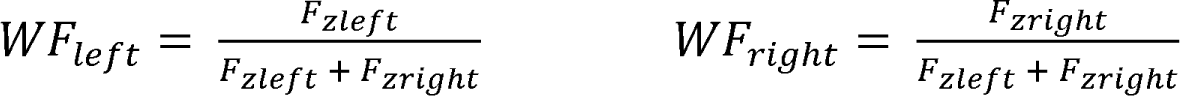

#### Touch forces

Touch force magnitude in z-axis (F_z_) and horizontal axis were measured directly from the touch plate readout.

#### Fluctuations

The means of the absolute values of the fluctuations were computed for CP_AP_, CP_ML_, WF and touch force fluctuations, for each trial.

#### Stabilogram Diffusion Function Analysis

Measures of postural balance show significant stochasticity and require analyses that do not implicitly filter out noise. One such analysis measures the mean square displacement (MSD) of any signal’s time series as a function of scanning time windows of varying widths. Stabilogram diffusion functions (SDF) introduced by Collins and De Luca (1993) to the study of balance are particularly sensitive to changes that occur in quiet stance tasks due to stochastic pattern changes that occur during different conditions, even when the sway amplitude averages may show relatively, stable, steady-state behavior across conditions (Balasubramaniam and Wing 2002, Bakshi, Ventura et al. 2014).

Stabilogram diffusion functions (SDF) were computed over the fluctuation time series: *C*(*t*_2_ − *t*_1_) = <[*q*(*t*_2_) − *q*(*t*_1_)]^2^>, where q represents a time-series of variables - CP_AP_, CP_ML_, WF and the touch force magnitudes as a function of time t.

#### Dependent variables and Statistical Analysis

Dependent variables derived from the fluctuations of WF and the AP and ML components of the CP were statistically analyzed in a 3-way MANOVA, within subject design, 3(Motion type) x 2(Touch and no touch) x 2(Order). The three levels of the motion type factor were Stationary scene, environment motion, and self-motion. The two levels of order of the task were “Start with motion followed by no motion” or “start with no motion followed by motion”.

Four dependent variables were also derived from the SDF analysis (following the equation C - Dr^a^^1^) of the fluctuations of CP_AP_, CP_ML_ and WF. These derived SDF parameters were i) the total area under each of the traces for different motion types, ii) the exponent a determined by the slope of log(C) plotted against log(r), iii) the diffusion coefficient (D) which measures the stochastic activity on average, and iv) the critical time point at which the power a - 2H - 1.

Both 2-way and 3-way interactions among the independent factors were analyzed with all four dependent variables. For pairwise comparison, we computed the contrast (type-repeated) between different levels of factors. Multivariate analysis incorporated Pillai’s Trace for significance at 0.05 level.

## Results

### Raw sample Data

Figure 1A and B provide raw sample data of net CP and WF, respectively, for one subject for four conditions. The left two panels are without touch, the right two are with touch. The top two are stationary to motion, the bottom two are the reverse. The central dashed black vertical line is when the transition between motion and stationary, or vice versa, occurs. The grey dashed vertical lines mark the onset times of the reported illusion of self-motion.

**Figure 1.**
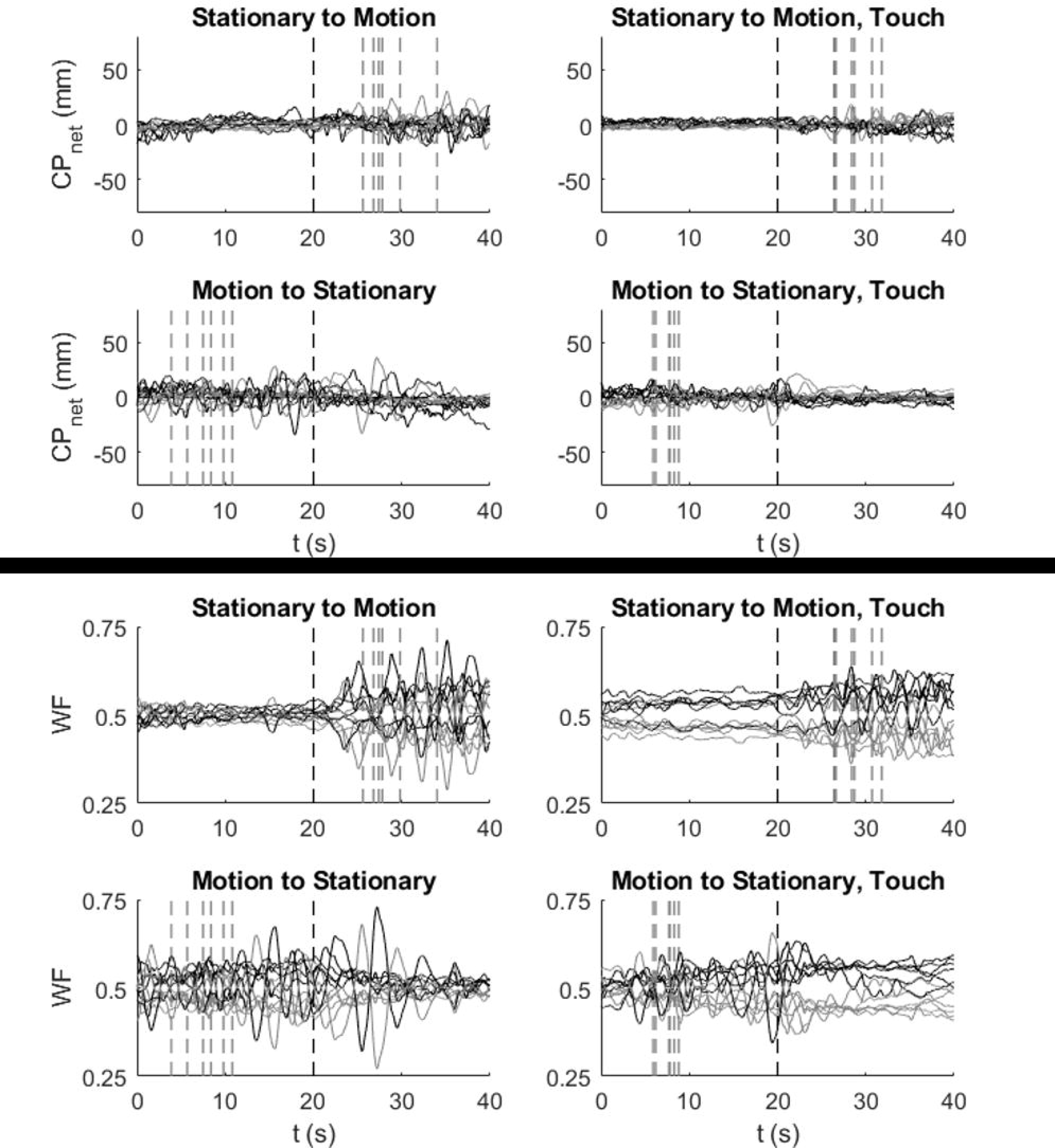
Sample data for a typical subject. The dark vertical dashed line at t=20 marks the change from stationary to motion (top panel) or vice versa (bottom). The grey vertical dashed lines mark the onset time of self-motion. **A)** CP_net_ traces across the four conditions (by motion order and touch). The AP component is in black and ML in grey. **B)** WF traces for the right (in black) and left (grey) foot. As the sum of WFs always equals one, the traces fluctuate symmetrically about the 50% horizontal level.

### Mean Fluctuations

A MANOVA testing for within-subjects effects for the three dependent variables (CP_AP,_ CP_ML_, and WF) was significant at <.05 level for the three motion-type levels (stationary, pre-, and per-self-motion illusion epochs, Pillai’s trace, F(6,50)=2.4, η^2^=.22). Touching also significantly stabilized balance, reducing the fluctuations by almost half (p<.01, F(3,11)=20.3, η =.85). There was no order effect – whether the subject’s faced the stationary condition first or after the rotation stimulus, it did not affect the average balance measure fluctuations.

In the AP direction, the perception of self-motion consistently gave higher mean net CP fluctuations (> 3 mm) than during perception of room-motion (< 3 mm). The univariates of CP_AP_ were significant for motion types (p<.01, F(2,26)=7.04, η^2^=.35) and touch (p<.01, F(1,13)=45.2, η =.78). For net CP fluctuation in the medial-lateral (ML) direction, there were significant differences arising from motion-type i.e. between different perception stages of the visual stimulation (p<.04, F(2,26)=3.69, η^2^=.22). Touch reduced the ML fluctuations mean significantly (p<.01, F(1,13)=51.5, η^2^=.80). The trend that self-motion gives higher fluctuation is consistent with that in the AP direction. An ANOVA for the fluctuations of the weight fraction (WF) in both feet across different conditions and different time phases shows a significant difference for motion type (p<.05, F(2,26)=3.4, η^2^=.21) and touch (p<.01, F(1,13)=50.5, η^2^=.79).

Note that the fluctuations in the left or right foot are absolutely equivalent, therefore, either one can be presented. The ANOVAs for the order effect was not interpreted for the three variables as the MANOVA was not significant. There was a significant two-way interaction between motion type and touch, for CP_AP_ (P<.04, F(2,26)=3.8, η^2^=.23), but not for CP_ML_ and WF.

In the post-hoc analysis, within-subject contrasts for motion-type show a significant difference between stationary and pre-self-motion for CP_AP_ (p<.01, F(1,13)=9.9, η^2^=.43), CP_ML_ (p<.01, F(1,13)=10.4, η^2^=.44), and WF (p<.01, F(1,13)=9.9, η^2^=.43). The contrast between pre- and per-self-motion was significant for CP_AP_ (p<.01, F(1,13)=12.8, η^2^=.49), CP_ML_ (p<.03, F(1,13)=6.1, η^2^=.32), and WF (p<.04, F(1,13)=5.7, η^2^=.31). The post-hoc within-subject contrasts for touch show significant difference between the two levels for all three dependent variables: CP_AP_ (p<.01, F(1,13)=45.2, η^2^=.78), CP_ML_ (p<.01, F(1,13)=51.5, η^2^=.80), and WF (p<.01, F(1,13)=50.5, η^2^=.79).

### SDFs of Fluctuations

For this analysis, the mean squared displacements were computed, as a function of different temporal windows widths, for the CP and WF fluctuations and the touch forces. We performed a three-factor MANOVA, with a 3 (Motion-type: stationary, environment moving, self-motion) X 2 (No-touch and touch) X 2 (Order of display stationary or moving) design, on the SDFs of the AP and ML component of CP fluctuations as well as the SDF of the WF fluctuations^2^. Four DVs were extracted from each SDF traces: Area, Diffusion Coefficient, Critical point, and Hurst Exponent). The measures derived from the SDFs showed significant main effects for touch, order and motion-type.

SDF of the anterior-posterior (AP) direction of the net CP are plotted, in Figure 2, against the temporal window width (τ). The SDF traces are presented for the three motion-type epochs: solid black line for when there was no motion displayed in the HMD; medium grey dashed line represents when subjects perceived environment rotation; and light grey dotted line is when subjects perceived self-motion. Each solid line trace is the average across all the subjects, and the corresponding shaded colored region represents the standard errors.

**Figure 2.**
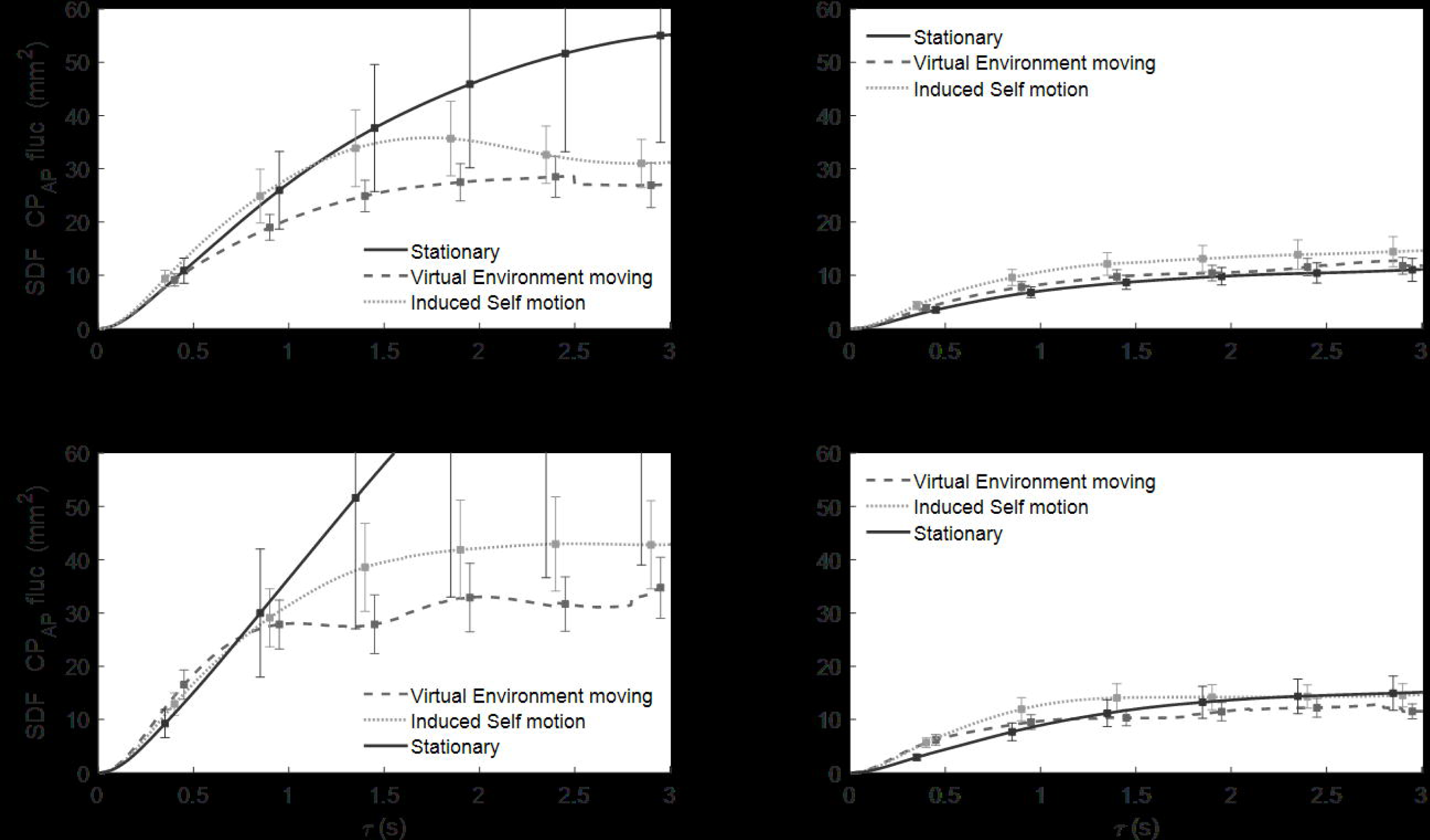
SDF vs. temporal width of the CP_AP_ fluctuations for all touch, motion-order conditions, and the three motion types.

No-touch trials are on the left panels, and touch trials are on the right. *No motion to motion* condition is presented in the upper subfigures, and *motion to no motion* is presented in the lower figures. Four distinct patterns emerge from the average SDF traces. For the no-touch conditions (left panels), the SDF trace of CP_AP_ fluctuations 1) during the stationary epoch is greater than all other motion types in the longer time scale (solid line above dashed or dotted lines), 2) during perceived self-motion is higher than that of perceived environment motion (dotted higher than dashed), 3) the stationary phase traces when preceded by motion (solid trace in the bottom-left panel) are significantly greater than when the trial starts with the stationary scene (solid black line in top-left). 4) The SDF traces of CP_AP_ fluctuations in the touch condition (right panels) are significantly smaller than those in no-touch conditions (left panel), demonstrating that non-supportive mechanical light touch overall reduces CP fluctuations significantly in all conditions despite any visual perceptual changes.

The CP_AP_ fluctuations showed significant main effects of motion type (F(8.000, 48.000) = 4.33, p < 0.01) and touch (F(4.000, 10.000) = 18.49, p < 0.001), but not for order. Univariates analysis showed that the significant difference in Motion-type arose from two of the DVs: the Hurst exponent (p<.001) and the Critical Coefficient (p<.02). The difference due to touch arose from three DVs, namely, Area (p<.01), Critical point (p<.01), and the Diffusion Coefficient (p<.01). Pairwise comparisons showed that the stationary versus the environment-moving levels of the motion-type factor were significantly different for the Hurst-exponent (p<.05) and Critical-point (p<.05), while self-motion was different from environment-rotation for the exponent (p<.05) only.

SDFs of CP fluctuations in the ML direction against τ are shown in Fig 3, where all line indicators and condition plots are the same as in Figure 2 above. In contrast to the CP_AP_ fluctuations SDF in Fig 2, when a trial started with no motion, SDF for CP_ML_ fluctuations were higher in perceived self-motion compared to that in stationary phase or perceived room rotation (dotted line higher than others in top-left panel). However, there was no difference between the SDFs of CP_ML_ fluctuations between stationary scene and perceived environment motion (solid and dashed lines in top-left). When the trial started with motion, SDF traces changed for different motion perceptions (bottom-left). The trials involving touch (right panels) were lower than no-touch (left panels) on average. The touch traces showed no significant difference among different perceptional states of motion, supporting previous evidence that light touch reduces CP_ML_ fluctuations significantly regardless of any visual perceptual changes.

**Figure 3.**
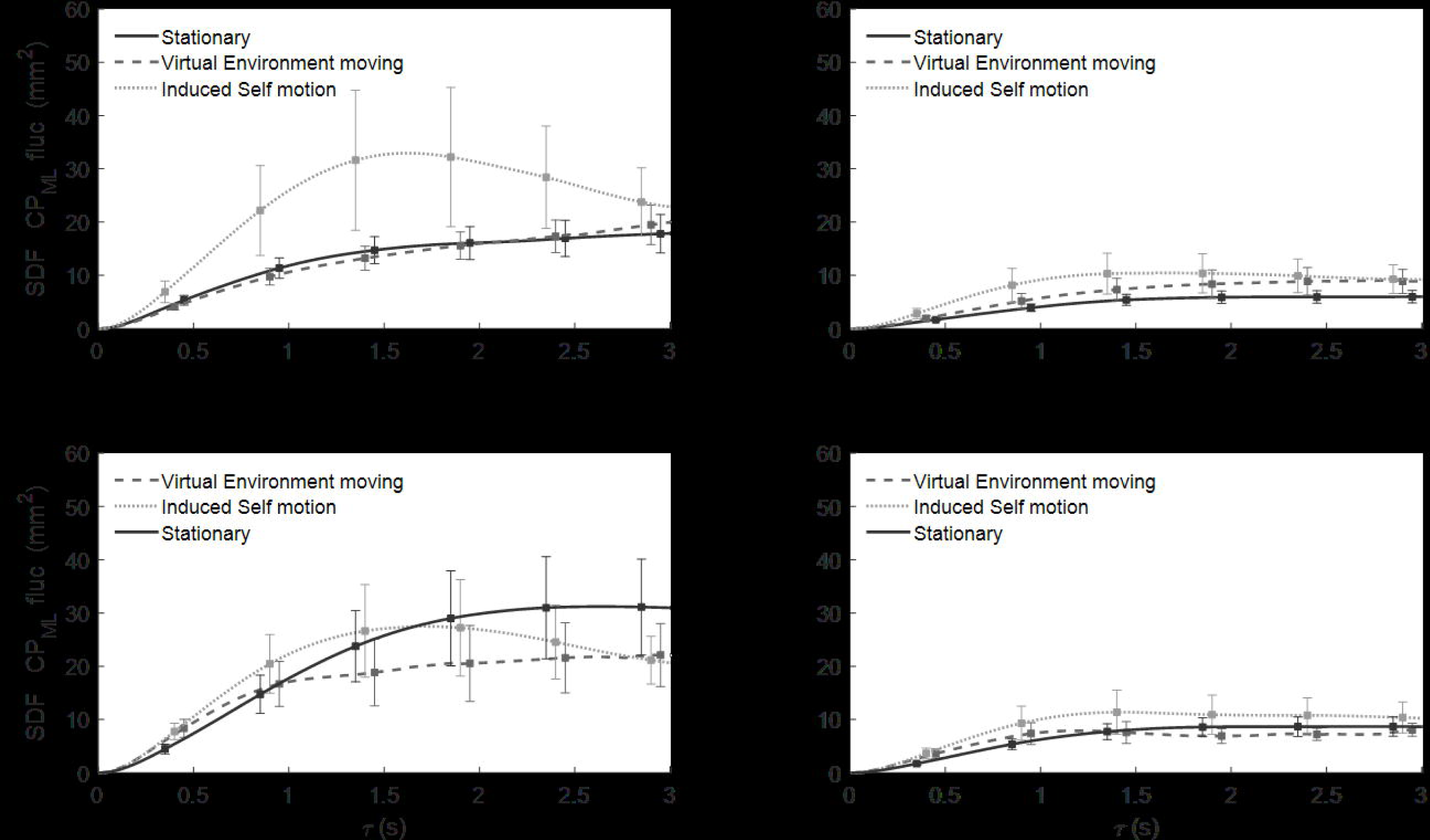
SDF vs. temporal width of the CP_ML_ fluctuations for all touch & motion-order conditions, and the three motion-types.

Statistically, the CP_ML_ fluctuations also showed significant main effect differences for motion-type (F(8.000, 48.000) = 4.99, p < 0.05) and touch (F(4.000, 10.000) = 6.85, p < 0.05), but not for order. The DVs that contributed to motion-type differences were the exponent (p<.001) and the critical point (p<.01), while only the area (p<.01) and diffusion coefficient (p<.001) had effects from touching vis-à-vis no-touching. Pairwise contrasts between the levels of motion type gave significant differences only for room-rotation versus stationary (p<.001 for Area and <.03 for Critical point), but not for self-motion.

SDF traces of WF fluctuation magnitudes in left foot (or right ^3^) are plotted against τ in Figure 4. All line indicators and condition plots are the same as in the two earlier figures. Similar to CP, WF fluctuations for touch trials (right panels) were reduced significantly vis-a-vis no-touch condition (left panels). In no-touch trials, when visual motion was provided after stationary scene, WF fluctuations during perceived self-motion (solid dark trace in top left) were significantly higher than those in no-motion and environment-motion phases. However, when the trials started with motion followed by no motion (bottom left panel), the no-motion trace was larger in average and variance, and there was no obvious dominance of any of the different phases of visual perception. The WF showed significant main-effects for all three factors – motion-type (F(8,48)=5.58, p<.001), touch (F(4,10)=7.63, p<.01) and order (F(4,10)=3.8, p<.05). The exponent and critical-point were the DVs that carried the significant differences (p<.001 and .01, respectively) for motion-type, while area and diffusion coefficient carried for touch (both <.01), but only diffusion-coefficient was significantly different for order. The stationary and the env. moving levels of motion-type were pairwise significantly different for the exponent (<.001) and the critical point (<.01). There was significant 2-way interaction between Motion-type and Order for all trials of fluctuations of CP_AP_ (p<.05), CP_ML_ (p<.01) and WF (p<.01). Motion-type also had a significant two-way interaction with touch for CP_AP_ (p<.05). The three factors also showed a three-way interaction for WF and CP_ML_ fluctuations (both p<.05), but not for CP_AP_.

**Figure 4.**
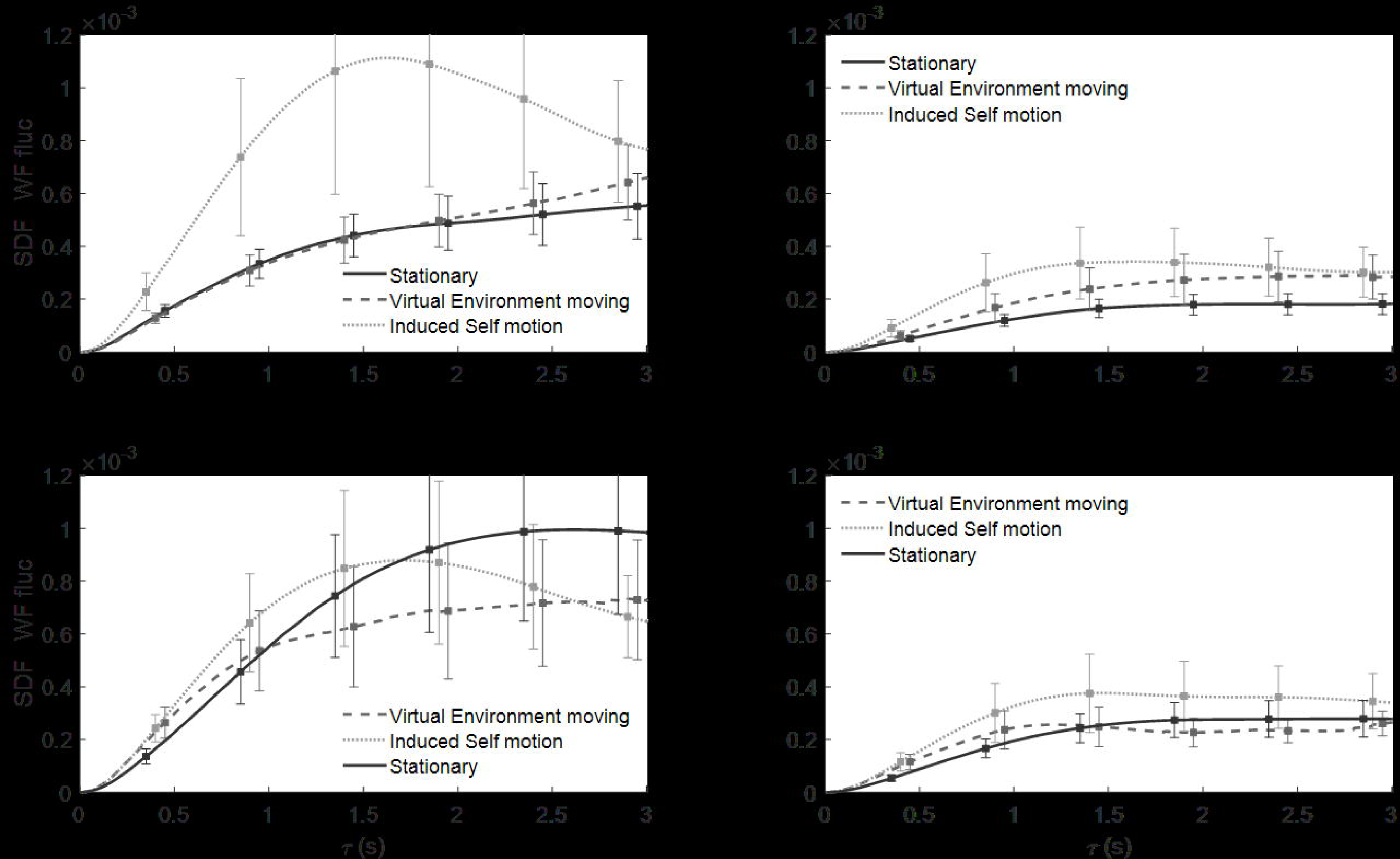
SDF of WF fluctuations.

### Touch Forces

The touch force in AP, ML, and Z axis, for the touch conditions, were analyzed for all order conditions and for the three different motion-type perceptual phases.

### There were no significant force magnitude differences with changes in the perceptual state of motion

The normal force component F_Z_ was around 40 gm. Incidentally, this is the magnitude at which the cutaneous receptors of the fingertips are most sensitive. The tangential components were relatively much smaller in magnitude: F_AP_ was between 5-10 gm, F_ML_ around 2g. Comparing the average amplitude of the fluctuations in the applied force magnitude during the touch trials, we found that AP forces were about 5 gms, ML was slightly lower, and the vertical forces were slightly larger. The touch magnitude fluctuation did not change significantly across motion epochs. Touch force fluctuations in the AP, ML and the Z-axis direction (normal to the ground) were measured, corresponding SDFs computed for each direction’s fluctuations. Touch force fluctuations in the AP and ML directions did not change significantly across conditions. When the trial was started with the stationary scene, the vertical force fluctuations showed no significant difference in touch force changes during different visual perception phases. However, when the trial started with motion followed by no motion, touch forces were greater when self-motion was perceived, and the effect continued-on during stationary phases. In comparison, exerted fingertip force fluctuations were lower during perceived environment motion.

### Latencies of onset of self-motion

There was no significant effect of touch on the time taken for the onset of self-motion. There was also no significant difference between touch and no touch trials on time taken for the onset of self-motion. However, there was a significant effect of motion order (F(1, 13)=15.8, p<.01), that is, there was a significant difference in report time whether the trial started with motion or started with no motion. The time of self-motion onset during stationary to motion condition (7.32s) was significantly larger than the time of self-motion onset during the motion to stationary condition (5.96s). Most subjects showed a report time within the average 5 to 10 seconds after motion started. The mean latencies of onset of self-motion were 7.6 ± 2.5s (stationary to motion without touch), 5.6 ± 2.1s (motion to stationary without touch), 7.1 ± 2s (stationary to motion with touch), and 6.3 ± 2.1s (motion to stationary with touch).

## Discussion

### Visual motion affects posture

It has been known, as discussed in the Introduction, that visual motion induces changes in postural sway. Our findings indicated a difference between environment rotation and self-rotation - in general, CP and WF fluctuation were higher during illusory self-motion than during the period when a moving visual scene was perceived. As our structured environment rotated in yaw, body sway under a quiet stance would be more perturbed in medial-lateral directions in the Stabilogram diffusion analysis. Our weight fraction analysis further showed that perceived self-motion considerably increased postural sway compared to that during perceived environment motion.

Our experiment elicited illusory self-motion using a virtual room’s 360° full-field rotation. Previous research typically has incorporated optic flow stimuli, moving dots or stripes, to induce a sense of self-motion and detect rotational velocity (Riecke 2011). Our task used a well-structured room environment that elicited both self-rotation and displacement, a sense of 360° self-motion. Normal optic flow provides important information by updating sensory inputs from self vs. environmental changes. Because our task involves a rotating visual environment at a constant speed, conflicting sensory inputs are provided over time, increasing postural sway instead of reducing it. For example, when self-motion is perceived, there are no angular acceleration signals from the semicircular canals to indicate motion onset. Typically, we consider self-motion perception to require integration of visual input, in the forms of optic flow, vestibular inputs that provide independent information of head movement, and proprioceptive as well as somatosensory information. In our study, no vestibular input was provided as the perception of self-motion was utterly illusory. Signals received by the visual cortex alone were strong enough to bring changes in postural balance task, reflected by a noticeable increase in weight loading fluctuations and the center of pressure fluctuations seen in the stochastic measures. When proprioceptive and somatosensory information were provided from the fingertip touch, they contradicted the visual information and reduced postural sway.

### Start with Motion vs. Start with No Motion

When the task began with optical rotation, and no touch cue was provided, moving visual input destabilized postural control more and led to a large aftereffect that persisted even when looking at a stationary scene later. It is worth noting that in all the statistical analyses for all variables, there was a main effect of motion type that is dependent upon the order of the motion-type – that is, different perceptions of motion depended on whether the trial began with motion or not. Motion aftereffects might explain this in the visual field (Andreeva 2016), where strong opposite visual motion would be induced after fast and short adaptation to a previously moving environment. In our task, while subjects’ eyes followed objects within the rotational environment, the sudden stop of motion might induce sudden eye movements in the direction opposite to the prior motion, after optokinetic nystagmus eye movements. This can create some level of motion sickness, as reported by some subjects and destabilize the maintained posture due to the long processing time in the sensory system to react to sudden environmental motion change. In future research, eye tracking will be employed to determine exactly how the eye moves during both the onset and stopping of visual motion.

### Touch vs. No Touch

In all test conditions, non-supportive mechanical fingertip touch significantly reduced the mean displacement of the center of pressure in both anterior-posterior and medial-lateral coordinates when visual input was introduced, as suggested by Figures 2 & 3. In particular, perceptual changes created by a moving visual input did not interact significantly with the effect of fingertip touch in maintaining postural balance, as touch generally reduced postural sway in all conditions. These results suggest that direct touch stabilization participates and belongs to a long-loop cortical reflex from the cutaneous receptors in the fingertip, possibly entrained in the parietal lobe and relayed to the primary motor cortex (Holden, Ventura et al. 1994). Out of 14 subjects, 9 subjects felt the touch plate moving with the fingertip when contact was present, 4 subjects did not sense the touch plate moving, and 1 subject sensed the touch plate switching from not moving to moving with the finger. However, overall, light touch reduced postural sway no matter what visual input was perceived; there were no significant differences among different SDFs of CP and WF for different motion types in touch conditions, which confirms a direct somatosensory and proprioceptive input of the touch entrained in postural sway. For the ten subjects, who experienced the touch plate moving with their finger, the motion of the tip of their finger and touch plate were precisely coordinative with their experienced body rotation, as if the plate were an extension of their own body.

In previous research, subjects’ voluntarily swaying in a slow rotating room had their sway deviated by Coriolis perturbations (Bakshi, DiZio et al. 2019). This postural sway was significantly attenuated by active light touch. In our research, postural sway generated during virtual environment rotation had a stochastic trajectory. Stabilogram measurements were a helpful tool in characterizing the changes in postural sway patterns that occur when only visual motion perception is altered. In combination, the fact that postural sway under both actual environment rotation and virtual environment rotation was significantly reduced by the addition of fingertip light touch suggests that touch reduces stochasticity of the task, probably independent of visual-vestibular integration for tasks involving the perception of self-motion.

### Touch magnitude

Consistent with our previous research, touch magnitude stayed at around 40 grams when visual motion was introduced. It did not change significantly with different phases of visual perceptual change, confirming that touch belongs to an independent long-loop reflex that adjusts postural sway using somatosensory and proprioceptive information.

### Author Contributions

All authors were involved in conceptualizing the experiment and contributed to the preparation of the manuscript. YY was the primary experimenter, recruiting subjects and running the experiment, and with AB’s help, wrote the first draft of the manuscript. AB was a secondary experimenter, and coded the data analysis scripts in MATLAB, prepared the figures, and did the statistical analysis in SPSS. All participated in interpretation of the results and the final manuscript.

## Footnotes

1 C is the stabilogram diffusion function which is the mean square displacement computed as the function of the temporal window width r. D is the diffusion coefficient, and a is related to the Hurst exponent.

2 Notably, the left or right foot WFs gave the same results, because knowing one also determines the other.

3 The output for the left and right foot fluctuations are exactly equivalent because WF_left_+WF_right_ =1; => Δ WF_left_ + Δ WF_right_ = 0; => |Δ WF_left_ | = |Δ WF_right_|. We plotted the weight fraction SDF for both feet separately in two figures and confirmed they were same.

